# Interactive Web-based Annotation of Plant MicroRNAs with iwa-miRNA

**DOI:** 10.1101/2020.12.01.405399

**Authors:** Ting Zhang, Jingjing Zhai, Xiaorong Zhang, Lei Ling, Menghan Li, Shang Xie, Minggui Song, Chuang Ma

## Abstract

MicroRNAs (miRNAs) are important regulators of gene expression. The large-scale detection and profiling of miRNAs has accelerated with the development of high-throughput small RNA sequencing (sRNA-Seq) techniques and bioinformatics tools. However, generating high-quality comprehensive miRNA annotations remains challenging, due to the intrinsic complexity of sRNA-Seq data and inherent limitations of existing miRNA predictions. Here, we present iwa-miRNA, a Galaxy-based framework that can facilitate miRNA annotation in plant species by combining computational analysis and manual curation. iwa-miRNA is specifically designed to generate a comprehensive list of miRNA candidates, bridging the gap between already annotated miRNAs provided by public miRNA databases and new predictions from sRNA-Seq datasets. It can also assist users to select promising miRNA candidates in an interactive mode through the automated and manual steps, contributing to the accessibility and reproducibility of genome-wide miRNA annotation. iwa-miRNA is user-friendly and can be easily deployed as a web application for researchers without programming experience. With flexible, interactive, and easy-to-use features, iwa-miRNA is a valuable tool for annotation of miRNAs in plant species with reference genomes. We illustrated the application of iwa-miRNA for miRNA annotation of plant species with varying complexity. The sources codes and web server of iwa-miRNA is freely accessible at: http://iwa-miRNA.omicstudio.cloud/.

## Introduction

MicroRNAs (miRNAs) are a class of small non-coding RNAs that are widespread in eukaryotes and play roles in a variety of biological processes, including plant growth, development, and stress responses [1–3]. In plants, miRNA genes are transcribed into primary transcripts, which are processed by the DICER-LIKE 1 (DCL1), SERRATE (SE), and HYPONASTIC LEAVES 1 (HYL1) proteins to generate a stem-loop structured miRNA precursor, following by trimming into a mature miRNA/miRNA* duplex [4]. Recently, some miRNAs have also been associated with agriculturally important traits, emerging as potential targets for crop improvement and protection [5]. Due to their biological and agricultural importance, miRNAs have become essential elements, annotated in genome sequences, particularly for plant species.

Genome-wide miRNA identification is generally accomplished using bioinformatics methods, such as homology search, machine learning-based prediction, and next-generation sequencing (NGS) data mining [6–14]. Such computationally identified miRNAs from multiple research groups have been deposited into public data repositories, such as miRBase [15], PmiREN [16], sRNAanno [17], and Plant small RNA genes (PsRNA) [18], providing valuable resources for life scientists interested in miRNA research, —from single-gene to genome-wide scale, basic molecular biology to population genetics, and bioinformatics to experimental biology. Despite these advances, present-day annotations remain riddled with false positives, and have a limited degree of comprehensiveness (the fraction of all *bona fide* miRNA genes that are included), exhaustiveness (the fraction of all mature miRNAs from each miRNA gene), and completeness (the fraction of pri/pre-miRNA sequences that cover the entire length) [19,20].

There are multiple factors that make the computational identification of miRNAs challenging. First, the tissue-/cell-type/developmental-stage specific expression and/or low expression properties of some miRNAs mean that they are often poorly identified from traditional low-throughput experimental studies and NGS experiments with limited samples and/or low sequencing depth. Second, the imperfect criteria defined for identification of miRNAs from NGS data. Although high-throughput criteria were established years ago [21], and have been continuously updated in response to studies of mechanisms [12,19], they cannot fully capture the species to species variation of miRNA characteristics. Third, the unsatisfactory level of accuracy of automatic miRNA annotation methods. Homology-based strategies fail to identify species-specific miRNAs, while machine learning-based tools have been designed for genome-wide miRNA prediction [7,8,22–24]; however, most were trained with limited experimentally validated miRNA data, and have markedly lower prediction accuracy in cross-species prediction [25]. Since the introduction of small RNA sequencing (sRNA-Seq), many sequencing-based tools have been developed that vary in their characterization of miRNAs [10–14]; however, only few tools have kept pace with updated miRNA identification criteria and they continue to suffer from trade-offs between quality and quantity [26]. Given these differences in the use of sRNA-Seq data, automatic annotation approaches, and miRNA identification criteria, inconsistency often arises in existing plant miRNA annotations. For example, at the beginning of this project in December 2019, we observed that, in *Arabidopsis thaliana,* there were 326, 221, 163, and 142 miRNA precursors annotated in miRBase (v22.1), PmiREN (v1.0), sRNAanno (v1.0), and PsRNA (v1.0) databases, respectively, with an overlap of only 120 miRNA precursors among these four databases. These inconsistencies indicate a proportion of false positives and false negatives within existing plant miRNA annotations, using which may result in unappreciated, but often serious, consequences for downstream studies, such as expression quantification, differential expression analysis, targetome analysis, and functional screening.

A straightforward way to improve the quality of miRNA annotations is to develop bioinformatics methods with sophisticated design of miRNA identification algorithms and criteria. In addition, a combination of automatic annotation and manual annotation would also be effective. The power of manual annotation has been demonstrated in human protein-coding gene annotation, where a team of human annotators inspect evidence supporting automatically annotated transcripts, and create relatively confident annotation databases, including GENCODE [27] and Ref-Seq [28]. These manual annotation databases are often free from many of the artefacts inherent in automated approaches, and have been adopted by most large-scale genomics projects, including the Encyclopedia of DNA Elements (ENCODE) [29] and the Genotype-Tissue Expression (GTEx) project [30]. In recent years, manual inspection has also been advocated, and performed to compile high-quality miRNA dataset from the genomes of human [31] and *Citrus sinensis* [26]. These pioneer investigations will provoke a wider interest among scientists in the research field of manual inspection of genome annotation, accompanied by an increased demand for effective interactive annotation tools to manage and analyze genome-wide miRNA annotations.

Here, we present iwa-miRNA, a web-based framework for interactive annotation of miRNAs from plant species with reference genomes. iwa-miRNA not only provides functions for automatically incorporating miRNA annotations from four representative databases (i.e., miRBase, PmiREN, sRNAanno, and PsRNA), but also builds a bioinformatics pipeline designed specifically to handle large-scale sRNA-Seq data for candidate miRNA prediction. Both annotated and predicted miRNAs are aggregated into a comprehensive list of miRNA candidates. Two miRNA selection approaches, high-throughput criteria and machine learning-based, are provided to assist the selection of promising miRNA candidates, based on the sequence-, structure-, and expression-based features. To enhance the accessibility of miRNA annotation, iwa-miRNA generates a report page with detailed information customized by feature types for each selected miRNA, facilitating convenient miRNA refinement during manual curation. The source codes of iwa-miRNA have been combined into a Galaxy platform, which are organized with a user-friendly web interface, and supported with extensive user documents. With these flexible, interactive, easy-to-use features, iwa-miRNA can generate a comprehensive collection of miRNA candidates and allows users to interrogate miRNA annotation in a straightforward way, without the need for computational skills. We provide examples of the application of iwa-miRNA for miRNA annotation of *Arabidopsis thaliana,* maize (*Zea mays* L.), and hexaploid bread wheat *(Triticum aestivum* L.).

## Method

The iwa-miRNA comprises three modules: ***MiRNA Compilation, MiRNA Selection,*** and ***Manual Curation*** (**Figure 1; Table 1**). The source codes of the modules and their dependencies are fully organized within the Galaxy framework. iwa-miRNA can be expertly implemented through a user-friendly web interface, and summarizes the results into HTML pages, using Rmarkdown for easy visualization, interpretation, and sharing.

**Figure 1.**
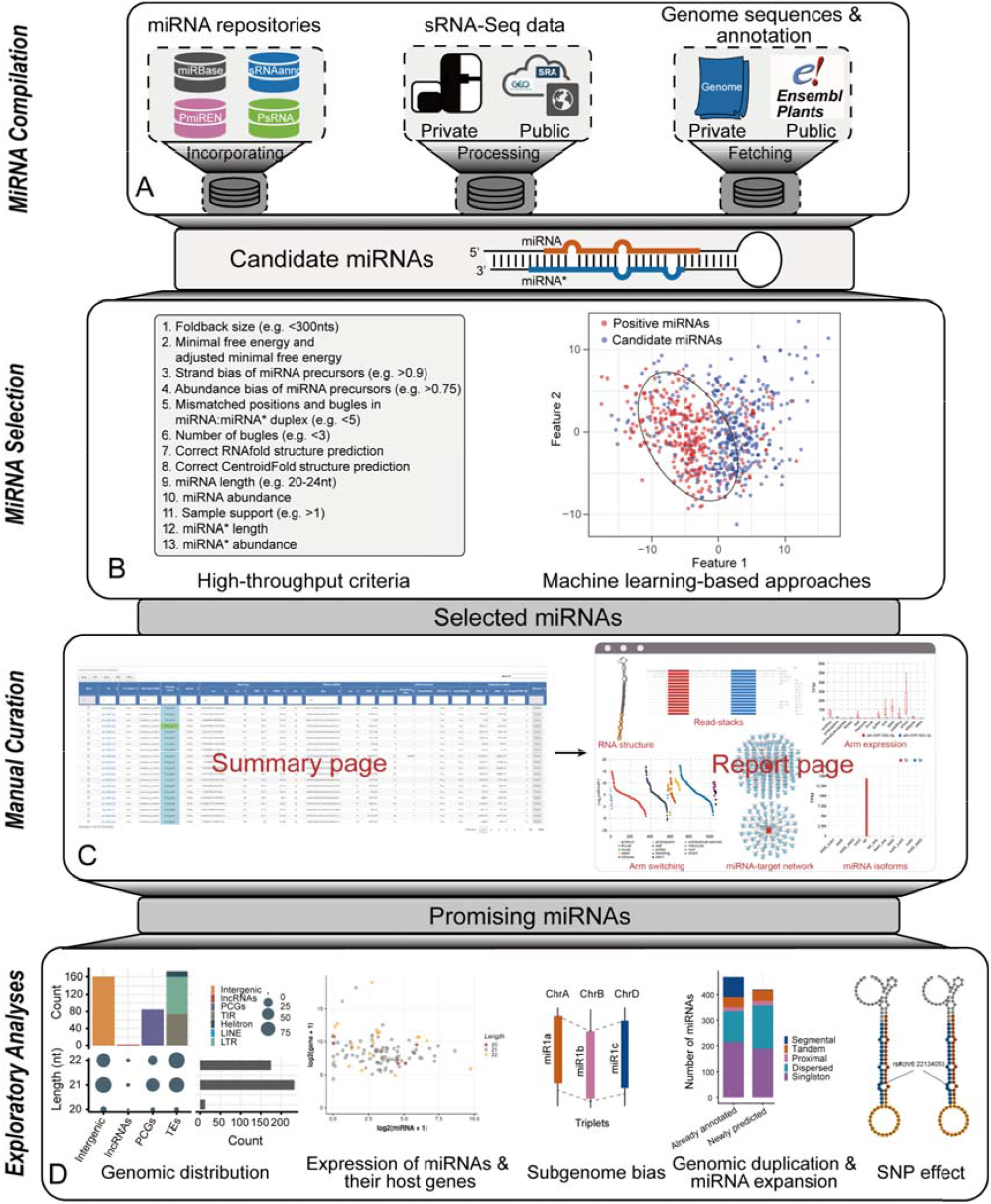
Graphical summary of iwa-miRNA. **A.** MiRNA candidates are generated by aggregating annotated and predicted miRNAs. B. Promising miRNAs are selected using high-throughput criteria and machine learning approaches. **C.** Manual curation of selected miRNAs based on annotation information from summary and report pages. **D.** Exploratory analysis of selected miRNAs. PCGs, protein coding genes; TEs, transposable elements; TIR, terminal inverted repeat; LINE, long interspersed element; LTR, long terminal repeat.

**Table 1.** Overview of functional modules in iwa-miRNA

| Module | Tool | Input | Output | Application |
| --- | --- | --- | --- | --- |
| MiRNA<br>Compilation | miRNAretrival | Name of species and miRNA databases | Already annotated miRNAs | Aggregate annotated miRNAs provided by four representative miRNA databases |
|  | genomePrepare | Name of species or genome sequences and annotation | Path of formatted genome sequences and annotation | Get genome sequences and annotation |
|  | miRNAPredict | SRA accession numbers or uploaded fastq files | Predicted miRNAs | Predict miRNAs from sRNA-Seq data |
|  | miRNATranslate | Output from miRNAretrival and miRNAPredict | miRNA and miRNA precursors with a uniform format | Translate annotated and predicted miRNAs into the genomic coordinate system |
| MiRNA<br>Selection | miRNASelection | Output from miRNATranslate | Selected miRNAs | Select promising miRNA candidates |
| Manual<br>Curation | manualCuration | Output from MiRNA Selection | Summary and report pages | Determine the quality of selected miRNAs |

***MiRNA Compilation*** (Module I): This module generates a comprehensive collection of miRNA candidates by aggregating already annotated miRNAs from four plant miRNA databases (i.e., miRBase, PmiREN, sRNAanno, and PsRNA) and predicted miRNAs from user-submitted sRNA-Seq data (Figure 1A). For a plant species of interest, all miRNA annotations (e.g. name, sequence, genomic coordinates, and so on) provided by the four miRNA databases are automatically retrieved using the “miRNARetrival” function. miRNA prediction is accomplished using the “miRNAPredict” function, which is specifically designed for parallel analysis of large-volume sRNA-Seq data. This function is based on a series of bioinformatics tools and custom scripts required for read cleaning (FASTX-Toolkit v0.0.14; http://hannonlab.cshl.edu/fastx_toolkit), genome mapping of reads (Bowtie v1.2.3 [32]), and miRNA prediction (miRDeep-P2 [12] and miRCat2 [33]). iwa-miRNA accepts the inputs of target genome sequences in FASTA format and corresponding annotations in GFF3/GTF format, which can be directly submitted by users or automatically fetched from the Ensembl Plants (https://plants.ensembl.org) database using the ‘genomePrepare’ tool. For miRNA annotations from different versions of the genome, the ‘miRNATranslate’ function can be used to translate annotated miRNAs into the genomic coordinate system of the target genome by performing miRNA precursor-to-genomic alignment using GMAP (v2019.09.12) [34]. All miRNA candidates are finally organized using a uniform naming scheme and genomic coordinates.

***MiRNA Selection*** (Module II): This module selects a subset of miRNA candidates that are regarded as promising miRNAs, according to the high-throughput criteria and/or using a machine learning-based approach (Figure 1B). For the latter miRNA selection approach, iwa-miRNA builds a one-class support vector machine (SVM) classifier [35] to predict if tested miRNA candidates are potentially real miRNAs or not (see File S1 and Table S1 for details). iwa-miRNA is user friendly, in that users can tune corresponding parameters according to the sRNA-Seq data at hand. A set of default parameters, derived from our own analysis experience, are also provided to assist non-expert users within their analyses.

***Manual Curation*** (Module III): This module provides the information for all miRNA candidates generated during the compilation and selection processes, and creates a summary page for rapid curation of the quality of selected miRNAs (Figure 1C). For miRNAs of interest, users can further inspect them by entering into the corresponding report pages, which provides more detailed information customized by feature types and visualized using various plot styles. A secondary structure plot is generated to display the location of a mature miRNA within the precursor sequence and quality-profiling results. Read stacks are plotted to show the read support of identified miRNAs. A boxplot is used to visualize miRNA expression patterns and arm selection events across different samples. A bipartite network is constructed to depict miRNA-target interactions predicted by psRNAtarget [36]. Users can update the list of selected miRNAs in a dynamic manner through adjusting criteria thresholds, or by direct deletion from the summary page. Selected miRNAs are finally exported into table, GFF3/GTF, and FASTA format files for downstream exploratory analysis (Figure 1D).

## Results

We illustrate the efficiency of iwa-miRNA for miRNA annotation of three plant genomes of different complexity. Among these applications, four databases (miRBase, PmiREN, sRNAanno, and PsRNA) and a set of publicly available sRNA-Seq datasets were used to generate candidate miRNAs. Both high-throughput criteria and one-class SVM with default parameters (Figure 1B) were used to identify promising miRNA candidates.

### Cases 1: Application of iwa-miRNA for miRNA annotation in *Arabidopsis*

As an initial demonstration of our framework, we looked at the long-studied and extensively annotated miRNAs of the model plant species, *Arabidopsis*, which has a relatively small genome of ~135 Mb. In *Arabidopsis,* miRNAs have been studied for over 18 years [37–39], and explored using more than 2000 sRNA-Seq datasets [40]. Using iwa-miRNA, we obtained a total of 365 miRNA precursors corresponding to 625 mature forms, from the four databases (miRBase, PmiREN, sRNAanno, and PsRNA; **Figure 2A**; Table S2). Using 1063 sRNA-Seq datasets (see details in File S1) for Columbia ecotype of *Arabidopsis* as inputs, iwa-miRNA predicted 435 miRNA precursors, 302 of which were not previously annotated in any of the four plant miRNA databases. This resulted in generation of 667 miRNA precursor candidates, corresponding to 1190 mature miRNA candidates for *Arabidopsis* (Table S2).

**Figure 2.**
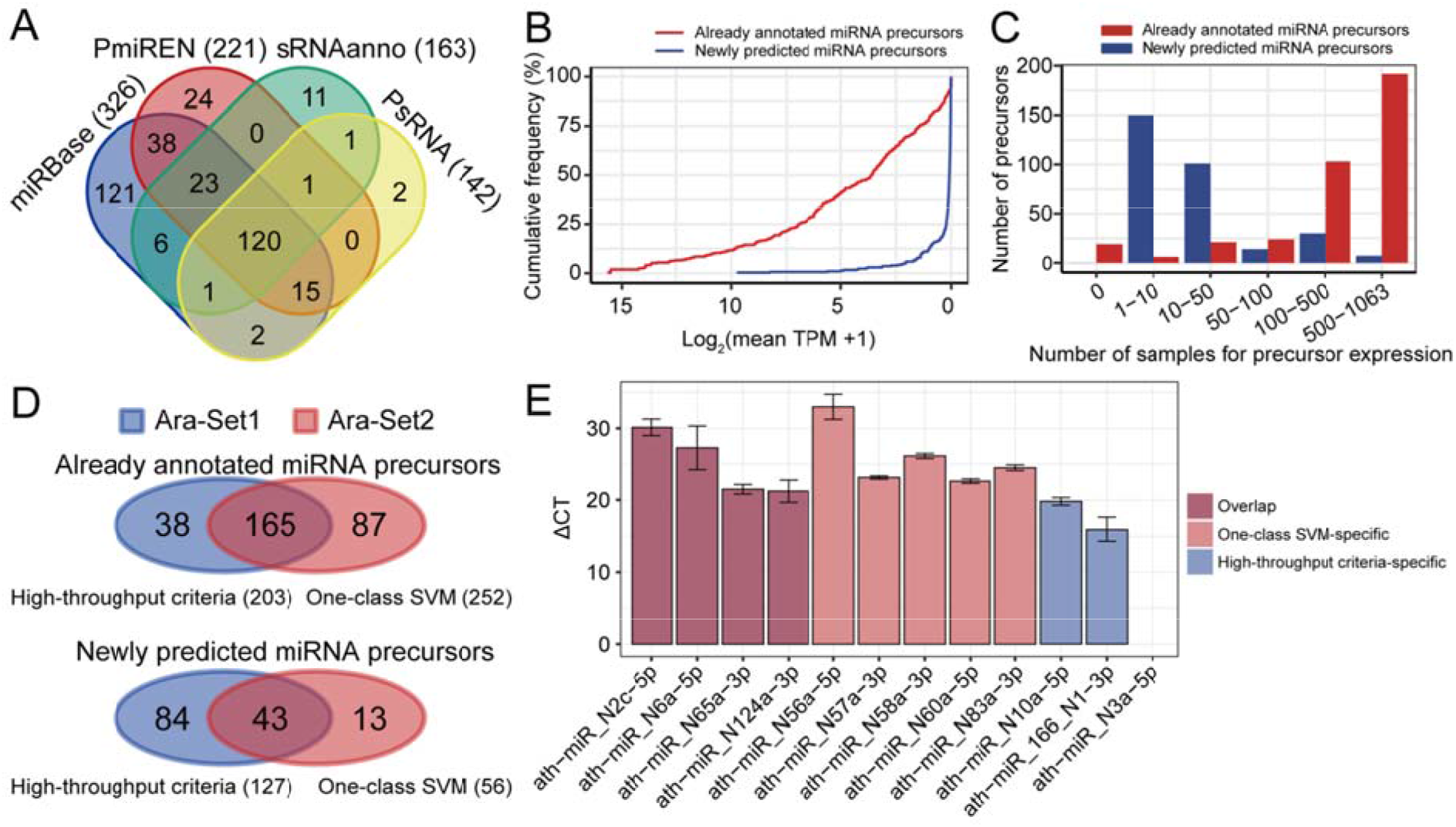
Application of miRNA annotation in *Arabidopsis*. **A.** Venn diagram comparing miRNA precursors provided by four miRNA databases. **B.** Cumulative frequency of log2 expression levels of already annotated and newly predicted miRNA precursor candidates. **C.** Number of expressed miRNA precursor candidates (already annotated and newly predicted) in different samples. **D.** Venn diagram comparing miRNA precursor candidates between the two miRNA selection approaches. **E.** Quantitative RT-PCR (qRT-PCR) results for 11 candidates. CT values were normalized to U6 small RNA. All values represent means ± SE (n = 3). TPM, transcripts per million; SVM, support vector machine; CT, cycle threshold.

Newly predicted miRNA precursors were expressed at remarkably lower expression levels and with less breath than those that were already annotated (Figure 2B–2C), indicating the potential importance of iwa-miRNA in identifying miRNA precursors with tissue-/cell-type/developmental-stage specific expression and/or low expression patterns. There were 330 miRNA precursor candidates that passed the high-throughput criteria (denoted as Ara-Set1), 203 of which were annotated in at least one of the four databases.

Using iwa-miRNA, we were able to characterize these 667 candidate miRNA precursors using 219 sequence features, 382 structural features, and 1,063 expression features (i.e., transcripts per million [TPM] values across all samples), providing an opportunity to predict miRNAs using machine learning approaches (see File S1 for details). Using 365 already annotated miRNA precursors as positive samples, iwa-miRNA built a one-class SVM classifier to predict 308 miRNA precursor candidates (Ara-Set2) as true positives. There were 208 candidate miRNA precursors in common between Ara-Set1 and Ara-Set2 (Figure 2D). For newly predicted miRNA precursors, 15 candidates (five from each region of the Venn diagram at the bottom of Figure 2D) were randomly selected for validation using quantitative real-time polymerase chain reaction (qRT-PCR) experiments, in which mature sequences were amplified with specifically designed primers (Table S3). Three miRNA precursor candidates were excluded because their mature sequences were not unique in the *Arabidopsis* reference genome sequence. These wet laboratory experiments validated 11 candidates as expressed in a mixed sample of *Arabidopsis* roots, shoots, leaves, and flowers (Figure 2E; File S1). These results provide evidence to confidently annotate *Arabidopsis* miRNAs using iwa-miRNA, although further validation experiments at large scale are desirable.

For each miRNA precursor candidate, iwa-miRNA assigns an identifier *via* a uniform naming scheme and the corresponding uniform resource identifier (URI) is hyperlinked to an HTML web page reporting a detailed description of feature information for manual inspection (**Figure 3**A). Figure 3B shows the report page of a representative example “ath-MIR156b”, which regulates vegetative phase change and recurring environmental stress by repressing squamosa promoter binding protein-like (SPL) transcription factors [41,42]. The precursor of miR156b produces two mature miRNAs of different lengths: a 20-nt miRNA from the 5’ arm (miR156b-5p) and a 23-nt miRNA from the 3’ arm (miR156b-3p). The former is high levels in root, leaf, and seed tissues, while the latter is preferentially expressed in root.

**Figure 3.**
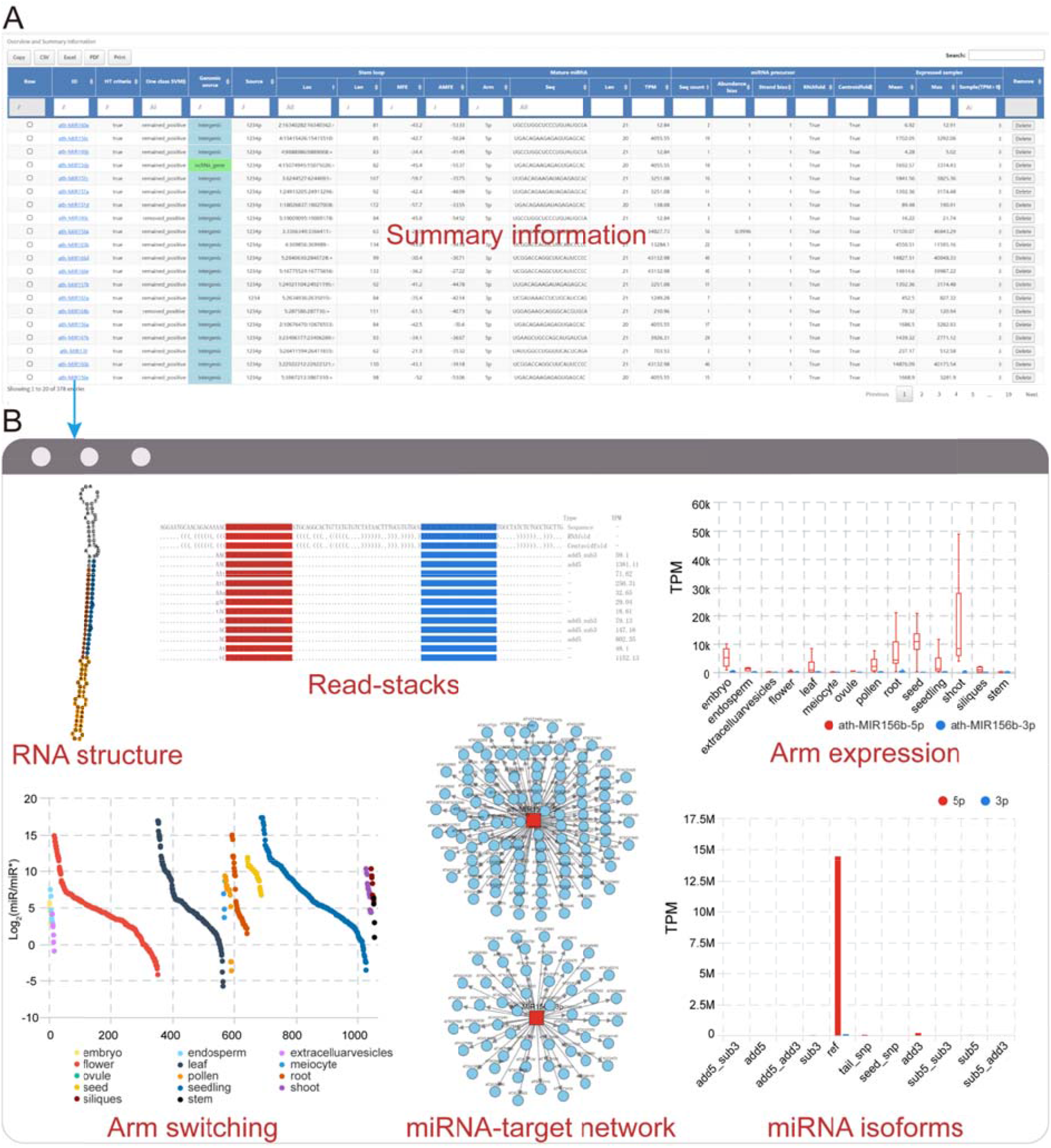
Summary and report pages generated by iwa-miRNA. **A.** Screenshot of summary page reporting the information of some features for *Arabidopsis* miRNAs. **B.** The report page for ath-MIR156b. HT, high-throughput; MFE, minimal free energy; AMFE, adjusted minimal free energy.

### Cases 2: Application of iwa-miRNA for miRNA annotation in maize

The successful application of iwa-miRNA to miRNA annotation in *Arabidopsis* prompted us to evaluate its value for analysis of plants with larger, more complex genomes. Here we focused on the model crop, maize, specifically the B73 inbred line, which has a reference genome of 2.3 Gb, more than 80% of which comprises transposable elements and other repeat sequences [43]. iwa-miRNA obtained a total of 619 miRNA precursors, which correspond to 893 mature forms, from the four databases (**Figure 4**A). For each database, the proportion of uniquely annotated maize miRNA precursors was markedly different from that observed in *Arabidopsis* (Table S4). This difference underscores the importance of performing an aggregation analysis and manual inspection of miRNA annotations from different sources. Further, on integrative analysis of 195 sRNA-seq datasets obtained from previously reported work [44], a total of 1241 miRNA precursor candidates were identified, 622 of which were previously un-annotated in any of the four plant miRNA databases (Table S5).

**Figure 4.**
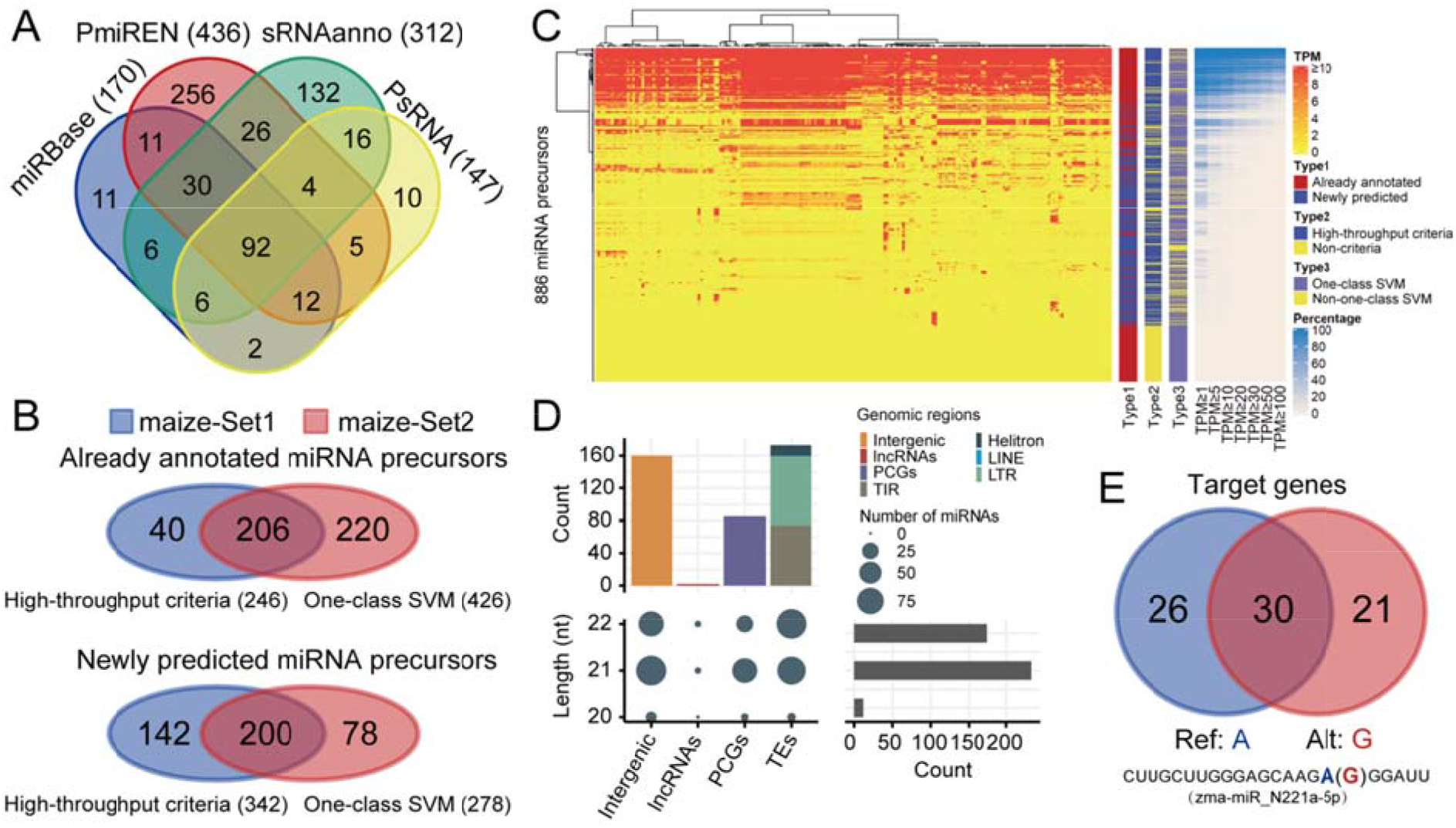
Application of miRNA annotation in maize. **A.** Venn diagram comparing miRNA precursors provided by four miRNA databases. **B.** Venn diagram comparing miRNA precursor candidates generated by two miRNA selection approaches. **C.** Expression levels of 886 miRNA precursors in 195 samples. Adjacent bar chart indicates three categorical results. The right panel shows the percentage of samples with expression greater than different thresholds (e.g. 1, 5, 10, etc.) **D.** Distribution of different miRNA lengths among different genomic features. **E.** The effect of SNP rs#chr6:22134053 on target genes of zma-MIR_N221a.

Using iwa-miRNA, 886 miRNA precursors were selected for downstream analysis: 588 passed with the high-throughput criteria (maize-Set1) and 704 predicted by the one-class SVM classifier (maize-Set2) (Figure 4B). One of the SVM-predicted miRNAs, zma-miR_N85a-5p, had already been validated by qRT-PCR in our recently published paper [44], and exhibited a specifical expression pattern in seed tissues. Of these 886 miRNA precursors, 381 exhibited broad expression patterns, having ≥ 1 TPM in more than 50% of 195 samples (Figure 4C).

Preliminary statistical analysis showed that most novel miRNAs were 21-nt and 22-nt in length (Table S5). Novel miRNAs were predominantly from intergenic regions and transposable elements (Figure 4D). Some newly predicted miRNAs are statistically associated with maize traits. As shown in Figure 4E, in the tail region of zma-miR_N221a-5p, there is a single nucleotide polymorphism (SNP; rs#chr6: 22134053) that has been reported to be significantly associated with the metabolic trait “Np.Npp_Feruloyl.caffeoyl_spermidine_derivative_E1” (*P*-value = 4.7E-6) [45]. This genetic variant (AA/GG) may influence target gene selection (56 *vs*. 51 genes; 30 overlapped). These preliminary results indicate the efficiency and power of interactive annotation of miRNAs in crop plants with complex genomes.

### Cases 3: Application of iwa-miRNA for miRNA annotation in wheat

Finally, we showcased the application of iwa-miRNA to miRNA annotation in a more complex crop plant, hexaploid (AABBDD) bread wheat *(Triticum aestivum),* which has an even larger genome (~17 Gb), with a more complex repeat landscape, than that of maize [46]. In addition to this, the wheat genome also presents other specific challenges, such as the composition of three closely-related and independently maintained subgenomes. Given that miRNAs were previously annotated based on different versions of the wheat reference genome from the four plant miRNA databases, iwa-miRNA first unified the miRNA annotation by mapping miRNA precursor sequences to the latest wheat reference genome (IWGSC RefSeq v1.0; https://plants.ensembl.org/Triticum_aestivum) using GMAP (v2019.09.12). Then, it was applied to predict miRNA precursors from 95 sRNA-Seq datasets (see details in File S1; Table S6), resulting in a list of 2857 miRNA precursor candidates in wheat (**Figure 5**A; Table S7). Subsequently, 2030 miRNA precursor candidates were selected, based on high-throughput criteria (wheat-Set1; 1617 miRNA precursor candidates) and one-class SVM prediction (wheat-Set2; 1410 miRNA precursor candidates) (Figure 5B). Finally, these 2030 selected miRNA precursors, corresponding to 2163 mature miRNAs, were organized into a summary page for future experimental validation and functional exploration.

**Figure 5.**
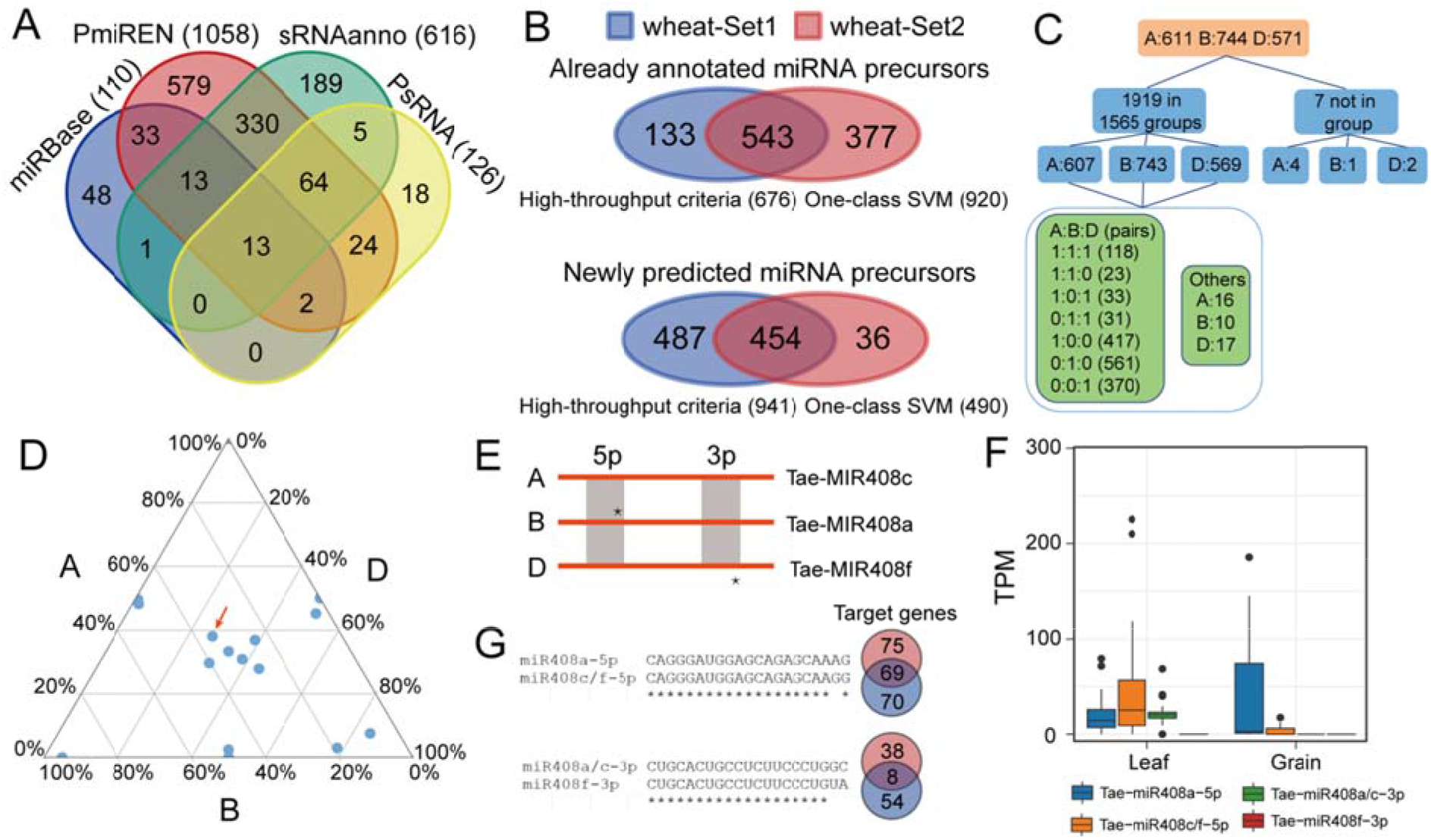
Application of miRNA annotation in wheat. **A.** Venn diagram comparing miRNA precursors provided by four miRNA databases. **B.** Venn diagram comparing miRNA precursor candidates generated by two miRNA selection approaches. **C.** Number of homologous groups and miRNAs in different A:B:D configurations. **D.** Ternary plot of miRNA expression levels from A, B, and D in triads. Tae-MIR408a/c/f is indicated by a red arrow. **E.** Schematic diagram of Tae-MIR408a/c/f in the ABD subgenomes. **F.** Expression levels of four mature sequences of Tae-MIR408a/c/f in leaf and grain tissues. **G.** The nucleotide difference and target alteration between mature sequences of Tae-MIR408a/c/f.

Of 2030 miRNA precursors, 1926 were clearly located on the three subgenomes, among which the D subgenome contained significantly fewer miRNA precursors than the A and B subgenomes (571 *vs*. 611 and 744, respectively; χ^2^ test, *P* < 0.001). This result suggests that there may be subgenome bias within these annotated miRNAs. Of 1926 miRNA precursors (A: 611; B: 744; D: 571), 1919 formed 1565 homologous groups of eight A:B:D configurations: 1:1:1 (118), 1:1:0 (23), 1:0:1 (33), 0:1:1 (31), 1:0:0 (417), 0:1:0 (561), 0:0:1 (370), and others (12) (Figure 5C; Table S8). Further, 7.54% (118/1565) of groups of homologous miRNA genes contained triads, with a single gene copy per subgenome. Among these 118 triads, 17 groups exhibited differences in mature sequences and expression levels (Figure 5D). A representative example is Tae-MIR408a/c/f, which has been linked to salt stress response and leaf polarity in wheat [5,47,48]. Among the A, B, and D subgenomes, there is a single nucleotide difference exists in the Tae-MIR408a/c/f 5’ mature sequence (A/B *vs*. D) and the 3’ mature sequence (A/D *vs*. B), resulting in identification of four mature sequences: Tae-miR408a-5p, Tae-miR408c/f-5p, Tae-miR408a/c-3p, and Tae-miR408f-3p (Figure 5E). These four sequences exhibit different expression patterns in leaf and grain tissues (Figure 5F). Tae-miR408a-5p and Tae-miR408c/f-5p were highly expressed in grain and leaf, respectively. Further, Tae-miR408a/c-3p was mainly expressed in leaf, while Tae-miR408f-3p had no expression in any tissues tested. A nucleotide difference in mature sequences also results in diversity in target genes (Figure 5G). These results indicate that the comprehensive annotation of miRNAs provides opportunities to explore evolution and functional diversification of miRNAs in polyploid plants, including hexaploid bread wheat.

## Discussion

MiRNAs have been the subject of extensive research over the past 20 years [39,49]; however, annotating miRNAs at the genome scale is not straightforward, not only because of the experimental difficulty in capturing some miRNAs with low- or context-specific expression patterns, but also owning to the computational difficulties in accurately distinguishing signals from noises within genomic sequences and sRNA-Seq data. To facilitate miRNA annotation, we have developed an interactive annotation framework, iwa-miRNA, with a user-friendly interface, to compile, select, and manually curate miRNAs from plant species with reference genomes.

Compared with existing miRNA-related bioinformatics annotation databases and tools, iwa-miRNA has several distinguishing features.

First, iwa-miRNA bridges the gap between existing annotations provided by public miRNA databases and new predictions from sRNA-Seq datasets. Most miRNA databases are periodically updated; however, they cannot integrate new miRNAs predicted from the rapidly accumulating sRNA-Seq data in a timely manner. In contrast, many bioinformatics tools have been designed with the sole purpose of predicting miRNAs from sRNA-Seq data, with less consideration of miRNA annotations provided by different databases. This issue was recently addressed by miRCarta [50], which collects novel human miRNA candidates and augments the annotation information provided by miRBase. Unlike miRCarta, iwa-miRNA tackles this deficiency with an emphasis on plant miRNAs. We suggest that, in the future, more attention should be paid to bridging the gap between miRNA annotations and predictions in plants, human, and other species.

Second, iwa-miRNA provides a new way to interactively select promising miRNAs. The aggregation of already annotated and newly predicted miRNAs generates a comprehensive collection of miRNA candidates, which certainly contain false positive hits, as well as interesting candidates (especially those expressed in tissue-/cell-type/developmental-stage specific patterns) for further validation. In previous studies, miRNA candidates were selected according to pre-defined rules, with ‘black box’ type functionality. The reliability of computationally annotated miRNAs is difficult to judge, particularly for scientists who are not involved in the process of miRNA prediction and annotation. iwa-miRNA provides a visualization of the sequence-, structure-, and expression-based features used in miRNA selection. In this way, with accessible miRNA annotation, researchers (both annotators and bench scientists) can manage miRNA selection in a dynamic manner, thus conveniently selecting promising candidates for further exploratory analysis and experimental validation.

Third, iwa-miRNA is user-friendly and can be deployed as a web application for easy accessibility and public/private data analysis. To facilitate the application of iwa-miRNA, all source codes and dependencies have been integrated into the Galaxy system, followed by packaging an independent runtime environment into a Docker image. This enables compatibility and portability: users can launch iwa-miRNA using a simple command, regardless of which operation system (Windows, Linux, or Macintosh), is used. Through a simple graphical interface, users can use this ‘one-stop’ systematic platform to mine massively accessible sRNA-Seq datasets and miRNA annotation. iwa-miRNA is also integrated with an Rmarkdown-based HTML report, to return interactive tables, publication-grade plots, and reproducible operations.

iwa-miRNA also suffers from some limitations. It cannot handle sRNA-Seq data from species without genome sequences. Recent RNA-Seq data analysis revealed that unmapped reads can serve as a valuable resource for new gene discovery [51]. In the current version, iwa-miRNA ignores sequencing reads that do not map to the reference genome. Since the main purpose of this study was to develop a platform that facilitates integrative annotation of miRNAs, iwa-miRNA has limited ability to perform downstream analysis of selected miRNAs, such as enrichment analysis (e.g. microRNA gene ontology annotations and miRNA set enrichment analysis) and comparative analysis between different species [52–54]. It should be also noted that there may be some false positives contained in the selected list of miRNA candidates. Further “wet” experiments should be performed to validate miRNA candidates of particular interest before any functional investigation.

## Future work

The iwa-miRNA project continues to be developed and improved. In future versions of iwa-miRNA, we plan to develop new functional modules to analyze sRNA-Seq data for species without a reference genome, identify candidate miRNAs from unmapped reads, and provide more downstream exploratory analysis. We invite researchers to use the iwa-miRNA platform to carry out large-scale sRNA-Seq data analysis on their local computers. Research collaborations are welcome, in particular for researchers without high-throughput computational resources.

## Data availability

iwa-miRNA Docker image is available at https://hub.docker.com/r/malab/iwa-mirna. Source codes and user manual are available at https://github.com/cma2015/iwa-miRNA. The prototype web server of iwa-miRNA is accessible at http://iwa-miRNA.omicstudio.cloud/.

## CRediT authorship contribution statement

**Ting Zhang**: Methodology, Software, Formal analysis, Data curation, Visualization, Writing – original draft, Writing – review & editing. **Jingjing Zhai**: Methodology, Software, Formal analysis, Writing – review & editing. **Xiaorong Zhang**: Data curation, Software. **Lei Ling**: Software, Formal analysis. **Menghan Li**: Data curation. **Shang Xie**: Software. **Minggui Song**: Software, Visualization. **Chuang Ma**: Conceptualization, Methodology, Supervision, Funding acquisition, Project administration, Writing – original draft, Writing – review & editing. All authors read and approved the final manuscript.

## Competing interests

The authors have declared no competing interests.

## Acknowledgements

We thank High-Performance Computing (HPC) of Northwest A&F University for providing computing resources. This work has been supported by the National Natural Science Foundation of China (31570371), the Youth 1000-Talent Program of China, the Hundred Talents Program of Shaanxi Province of China, and the Fundamental Research Funds for the Central Universities (2452020041).

## Supplementary material

**File S1 Detailed information regarding sRNA-Seq data collection and preprocess, SVM modeling, qRT-PCR experiment, and syntenic analysis of the wheat genome**

**Table S1 Sequence and structural features used in iwa-miRNA**

**Table S2 List of candidate miRNAs and their precursors in *Arabidopsis***

**Table S3 Primers for miRNA qRT-PCR assay in *Arabidopsis***

**Table S4 Proportion of miRNA precursors uniquely annotated in each of four miRNA databases**

**Table S5 List of candidate miRNAs and their precursors in maize**

**Table S6 List of 95 wheat sRNA-Seq libraries**

**Table S7 List of candidate miRNAs and their precursors in wheat**

**Table S8 Homologous groups of A:B:D configurations and detailed miRNAs in wheat**

